# Automated, image-based quantification of peroxisome characteristics with *perox-per-cell*

**DOI:** 10.1101/2024.04.08.588597

**Authors:** Maxwell L. Neal, Nandini Shukla, Fred D. Mast, Jean-Claude Farré, Therese M. Pacio, Katelyn E. Raney-Plourde, Sumedh Prasad, Suresh Subramani, John D. Aitchison

**Affiliations:** Seattle Children’s Research Institute, Center for Global Infectious Disease Research, Seattle, WA, USA; Department of Molecular Biology, School of Biological Sciences, University of California, San Diego, La Jolla, CA, USA

**Author notes:** To whom correspondence should be addressed. Correspondence to: Maxwell L. Neal, Ph.D., Senior Scientist, Center for Global Infectious Disease Research, Seattle Children’s Research Institute, 307 Westlake Avenue North, Suite 500, Seattle Washington, 98109-5219, OFFICE.

## Abstract

*perox-per-cell* automates cumbersome, image-based data collection tasks often encountered in peroxisome research. The software processes microscopy images to quantify peroxisome features in yeast cells. It uses off-the-shelf image processing tools to automatically segment cells and peroxisomes and then outputs quantitative metrics including peroxisome counts per cell and spatial areas. In validation tests, we found that *perox-per-cell* output agrees well with manually-quantified peroxisomal counts and cell instances, thereby enabling high-throughput quantification of peroxisomal characteristics. The software is available at https://github.com/AitchisonLab/perox-per-cell

## INTRODUCTION

Peroxisomes are membrane-bound organelles found in all eukaryotic cells. Originally defined based on their production of hydrogen peroxide from oxidative reactions, peroxisomes perform various functions including fatty acid oxidation, lipid biosynthesis, and reactive oxygen species metabolism. Peroxisomes are produced in cells either through *de novo* biogenesis, where the organelle is assembled anew, or through growth and division. In humans, mutations in genes encoding proteins required for peroxisome biogenesis genes cause severe developmental defects, collectively referred to as the Zellweger Spectrum of Peroxisome Biogenesis disorders. While some of the molecular mechanisms underlying peroxisome biogenesis have been uncovered, our understanding of the process remains incomplete (Mast *et al*., 2020; Farré *et al*., 2019). Motivated by a need for high-throughput analyses to reveal critical components contributing to this process, we developed open-source software called *perox-per-cell* for image-based quantification of peroxisome characteristics in yeast cells. The software automates several time-consuming, traditionally manual image processing tasks including the segmentation of yeast cells and their peroxisomes as well as the collection of quantitative metrics such as the number of peroxisomes in individual cells, the spatial areas of cells and peroxisomes, and the luminal signal intensity within segmented peroxisomes. Through this automation, we have substantially increased the speed at which we can assess and quantify the impact of cellular and genetic perturbations on peroxisome number and size. Here, we detail the methods underlying *perox-per-cell* and demonstrate its validity for accurately quantifying metrics relevant for peroxisome research.

## METHODS

*perox-per-cell* was developed to analyze microscopy data collected from yeast cells stained with calcofluor white demarcating cell boundaries and producing GFP tagged with peroxisome targeting sequence 1 (PTS1) to locate peroxisomes. The software enables rapid assessment of peroxisomal phenotypes from 3D-imaging Z-stacks of the calcofluor and GFP signals captured at the same physical position. With *perox-per-cell* we aimed to improve on previous work (Niemisto *et al*., 2006) and create a rigorously-tested, publicly available tool for the research community. *perox-per-cell* first projects the Z-stacks onto a single 2D image plane for both the calcofluor and GFP channels and then segments the cells and peroxisomes using automated tools. It then assigns peroxisomes to individual cells based on the overlap between the cell and peroxisome segmentation masks (Figure 1A-B). Finally, the number of peroxisomes in each cell and additional metrics are output to an Excel spreadsheet.

**Fig. 1.**
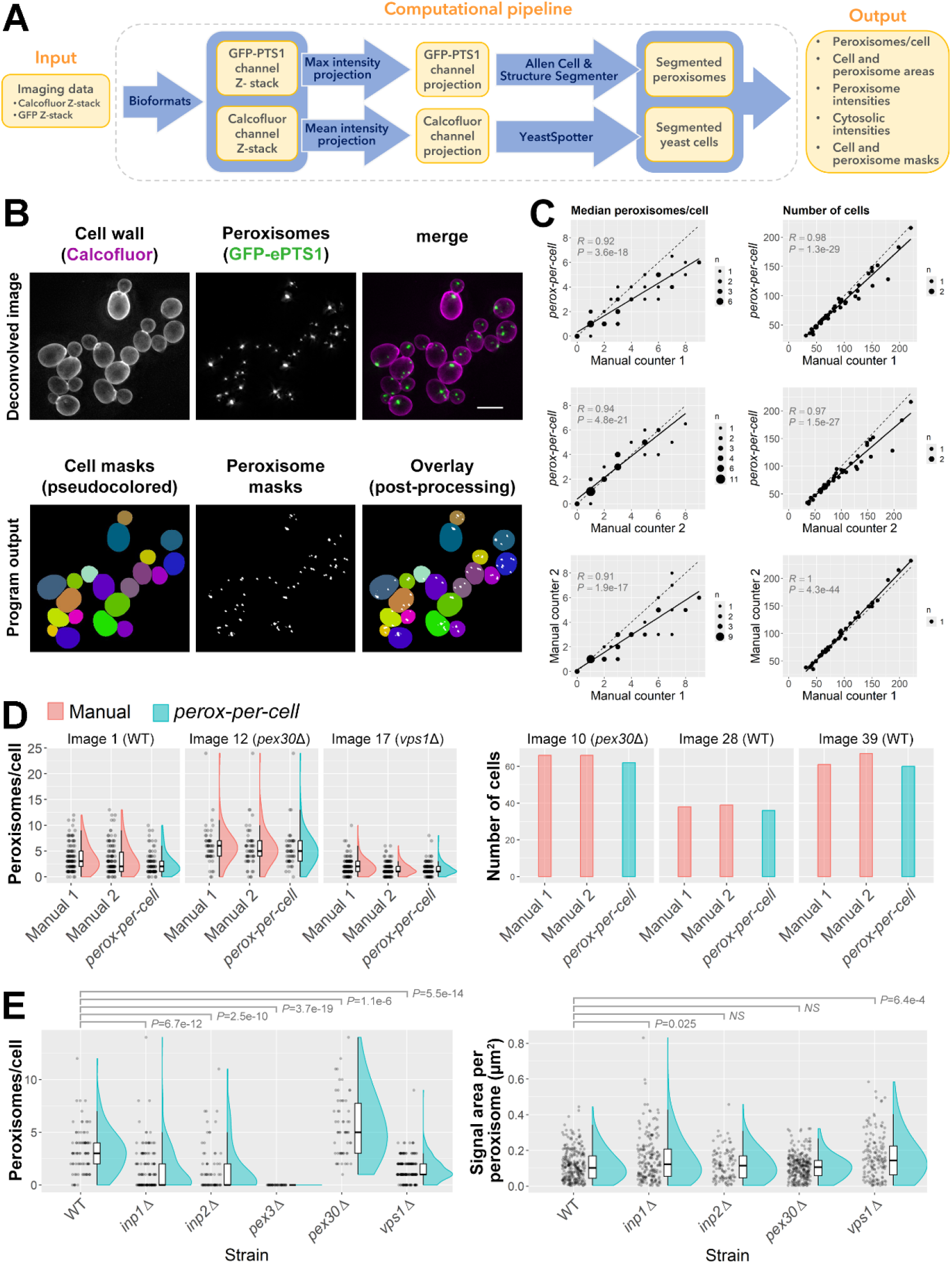
A) Architecture of the *perox-per-cell* computational pipeline. B) Example comparison of deconvolved images (top) used for manual counting and the cell and peroxisome masks generated by *perox-per-cell* (bottom). Scale bar: 5 μm. C) Scatterplots with trendlines (solid) and unity lines (dashed) showing correlation between manual and auto-generated peroxisome counts (left) and number of cells (right) across test images. *R*: Pearson’s correlation coefficient; *P*: correlation *P*-value; n: number of observations at coordinateøs. D) Distribution of peroxisome counts (left) and number of cells (right) derived manually or via *perox-per-cell* for three images where agreement between the methods was most representative of the overall agreement across test images. Distributions are shown using adjacent jitter, box, and half-violin plots. E) Distribution of peroxisome counts (left) and areas (right) from images most representative of a given strain’s peroxisomal features across test images. FDR-adjusted *P*-values from Wilcoxon rank-sum tests comparing mutant strains to WT are shown above plots.

### Z-stack intensity projections

Imaging files are read into *perox-per-cell* using Bio-Formats (https://github.com/CellProfiler/python-bioformats). *perox-per-cell* uses file metadata to initialize 3D arrays of appropriate dimension to store the Z-stack intensity values in the calcofluor and GFP channels. The program is designed to process input data consisting of two imaging channels, one for segmenting peroxisomes, the other for cells. After reading in the Z-stacks, a 2D maximum intensity projection is generated from the peroxisome channel Z-stack to ensure that each peroxisome, regardless of its position in the Z-dimension, is represented. The cell channel Z-stack is also converted into a 2D image so that it can be processed by the cell segmenter. For the cell channel, we use an *average* intensity projection because this resulted in more accurate segmentations in initial tests. Such projections may de-emphasize the presence of high intensity calcofluor signals localizing on bud scars, generating a more even intensity profile along cell boundaries. We note that *perox-per-cell* also accepts 2D images; a full 3D Z-stack is not required for either channel.

### Cell segmentation

Several machine learning (ML) tools for automatic segmentation of yeast cells have become available in recent years (Lu *et al*., 2019; Salem *et al*., 2021; Dietler *et al*., 2020). Based on visual inspection, applying the computational methods used by the YeastSpotter tool (Lu *et al*., 2019), which segments 2D images using a region-based convolutional neural network, gave sufficiently accurate segmentations. We initially chose YeastSpotter because it outperformed other established methods for segmenting fluorescently-imaged cells (Lu *et al*., 2019). We also tested YeastNet (Salem *et al*., 2021); however, it was trained on brightfield images and did not identify any cells when applied to test images. To implement YeastSpotter segmentation, we adapted the code at https://github.com/alexxijielu/yeast_segmentation, only making a minor change to ensure our tool loads the underlying model weights correctly when deployed as a standalone application. After performing cell segmentation on the calcofluor channel 2D projection, *perox-per-cell* saves the resulting image mask where all pixels corresponding to an individual cell share a unique intensity value.

### Peroxisome segmentation

To segment peroxisomes, we adapted one of a suite of scripts in the Allen Cell & Structure Segmenter package (https://www.allencell.org/segmenter.html), which provides workflows for automated segmentation of various subcellular structures. After exploring example workflows in the imaging software Napari, we adapted the “gja1” workflow, which segments punctate features. Our workflow performs Gaussian smoothing on the 2D projection from the peroxisomal imaging Z-stack followed by spot detection via Laplacian of Gaussian filtering. Spot detection sensitivity is set via a user-defined input parameter that can be adjusted if too many, or too few, puncta are being detected as peroxisomes. By default, this parameter is set to a value that results in reasonable peroxisome segmentation based on initial test images of various yeast strains. An additional user-specified parameter sets the minimum peroxisome size, in pixels. Segmented peroxisomes below this size, which specifies the limit below which peroxisomes cannot be identified with high confidence, are removed from downstream analysis. Following these steps, a binary mask of the segmented peroxisomes is output as well as a mask where pixels within individual peroxisomes share a unique label.

### Quantification of peroxisome abundance and other metrics

To quantify the number of peroxisomes in each cell, *perox-per-cell* iterates over individual cells in the cell segmentation mask and counts the number of peroxisomes in the peroxisome segmentation mask found within each cell’s boundary. Cells with pixels on the image border are excluded because their full areas may not be represented, leading to some undercounting. Each individual peroxisome is assigned to only one cell. If a peroxisome overlaps with multiple cells, it is assigned to the cell with the greatest overlap. In cases where a peroxisome’s maximum overlap is the same across multiple candidates, the peroxisome is assigned to the cell containing the greater signal intensity from the peroxisome channel projection image. If still unassigned after these steps, the peroxisome is randomly assigned to one of the candidates. The number of peroxisomes in each cell is tabulated and then output to a spreadsheet.

*perox-per-cell* also generates metrics for peroxisomal phenotypes, including the physical areas of each cell and peroxisome, the total summed intensity of the peroxisomal channel within cell boundaries, and the same summed intensity excluding signals from segmented peroxisomes. Accurate quantification of cell and peroxisomal physical areas requires that the physical size of pixels be specified in the input image metadata. To compute physical areas, *perox-per-cell* reads in metadata formatted according to the OME-XML standard (Goldberg *et al*., 2005) embedded in the input image. To accurately quantify areas, users should ensure that pixel width, height and associated units are included in that metadata.

### Software implementation, availability, and execution

The software is open-source, implemented in Python, and available at https://github.com/AitchisonLab/perox-per-cell. We have also deployed *perox-per-cell* as a standalone, Windows executable where users edit a batch file to set the location of their input imaging data and can adjust analysis parameters controlling peroxisome detection sensitivity, minimum peroxisome area, and the maximum possible intensity value in the peroxisomal channel. This last parameter should be set based on the bit depth of the peroxisome channel imaging data, equivalent to 2^(bit depth)^ – 1.

## RESULTS

To evaluate *perox-per-cell*, we compared its performance against manually-quantified data. We collected 44 two-channel, 14-bit Z-stacks from various yeast strains including WT as well as *inp1*Δ, *inp2*Δ, *pex30*Δ, and *vps1*Δ mutants. These strains were selected because, together, they encompass a variety of peroxisomal densities and cellular distributions among strains relevant to peroxisome research. For example, *vps1*Δ cells predominantly have a single peroxisome whereas *pex30*Δ mutants show elevated peroxisome counts (Hoepfner *et al*., 2001; Vizeacoumar *et al*., 2003). Each imaging data set consisted of a Z-stack of calcofluor-stained cells and a Z-stack from the same position to image GFP-PTS1 (see Supplemental Methods). For manual counting, stacks were first deconvolved and then converted to maximum-intensity projections. Two individuals manually counted the number of peroxisomes in each cell from these images using blinded samples. We then ran *perox-per-cell* on the same imaging data and compared the results against manual counts.

### Validation against manual peroxisome and cell counts

*perox-per-cell* results correlated strongly with manual counts (Figure 1C, left; Figure S1). On average, the median number of peroxisomes per cell estimated by *perox-per-cell* in an image differed by -0.44 peroxisomes compared to the average of the two manual counters (Figure 1D, left). Comparing *perox-per-cell* to manual counters 1 and 2 independently, this difference was - 0.91 and 0.03, respectively. The difference between manual counters was 0.94, illustrating that *perox-per-cell* estimates fall within the variability that exists between manual counters. We also assessed the agreement between *perox-per-cell* and manual counts in terms of cell-to-cell variability. The average difference between the interquartile ranges (IQRs) of *perox-per-cell* and manual peroxisome counts was -0.33 across images. The IQR from manual counter 1 was 0.45 peroxisomes per cell higher than counter 2 on average, also indicating that software estimates fall within manual counter variability.

*perox-per-cell* also showed strong correlation with manual cell number counts (Figure 1C, right; Figure S2). Across images, *perox-per-cell* detected a median value of 6.4% fewer cells (IQR = 6.5%) when compared to average counts recorded manually (Figure 1D, right). Inspecting images with the largest discrepancies between manual and automated counts, we found that cells not detected by *perox-per-cell* tended to have lower signal intensities and were crowded by neighboring cells. We have also noticed that the cell segmentation tends to not detect very small buds, possibly due to relatively lower calcofluor staining and, thus, lower signal intensity. We therefore encourage users to ensure that cell staining is sufficiently strong, and that cell crowding is minimal in input images.

### Detecting altered peroxisome characteristics in mutants

We investigated whether the results from *perox-per-cell* could be used to detect statistically significant alterations in cellular peroxisome counts among mutant strains with peroxisomal defects. For each strain represented in our set of 44 test images, we selected a representative image for each strain (Supplemental Methods), then compared peroxisome count distributions in mutant strains to WT. As part of this analysis, we also included a set of four images from a *pex3*Δ strain (Höhfeld *et al*., 1991) in which no peroxisomes were visually detected. We found that for all mutant strains tested, statistical comparisons against WT were significant with FDR-adjusted *P*-values ≤ 1.1e-06 for each strain (Figure 1E, left).

We also tested whether peroxisome areas differed between strains and found that *inp1*Δ and *vps1*Δ had significantly higher areas compared to WT (Figure 1E, right). We note that using the GFP channel intensities for this quantification is a proxy for peroxisomal area; differences in area could also represent differential PTS1 import. Together, these results indicate that *perox-per-cell* results can reveal statistically significant alterations in peroxisome characteristics in strains with known defects.

## DISCUSSION

Our results show that *perox-per-cell* accurately quantifies peroxisome counts from imaging data and that its estimates fall within the variability between manual counters. On a standard Windows workstation, the program processes images such as those used here in less than 90 seconds. Therefore, we anticipate it will be useful for rapidly and accurately assessing peroxisome characteristics in yeast cells, which, as we show here, can reveal statistically significant differences in peroxisome phenotypes between strains using one imaging instance per strain (Figure 1E).

While calcofluor staining was used to delineate cell boundaries in data used for this study, other staining methods may be applicable for use with *perox-per-cell*. The ML model used for cell segmentation was trained on fluorescent cell nuclei images, and additional studies that examine its generalizability across other staining methods and image modalities are warranted. Furthermore, the peroxisome segmentation method is not specific for peroxisomes and could be used for segmenting other types of intracellular puncta, provided that the user-defined parameters for peroxisome segmentation (see Methods) are tuned accordingly.

With *perox-per-cell*, we aim to facilitate peroxisome research in the domain of big data and its attendant analytical methods. The software opens new vistas for statistically-powered investigations that establish or refine definitions of peroxisomal defects and that uncover novel phenotypes of known mutants unresolvable through coarse-grained analyses.

## ACKNOWLEDGEMENTS

We thank members of the Subramani and Aitchison labs for useful discussions.

