## Supplemental Methods for "Automated, image-based quantification of peroxisome characteristics with *perox-per-cell*"

### Supplementary Data

Contents


- Supplementary Methods
- Table S1
- Figure S1
- Figure S2


---

#### Supplementary Methods

##### *Image acquisition*

For all experiments, cells were grown in synthetic defined medium (SD: 6.7 g/L Yeast nitrogen base without amino acids + 0.79 g/L CSM) with 2% Dextrose in flask cultures shaken at 250 rpm at 30 °C until log phase after which they were pelleted and resuspended in 50 µg/ml calcofluor white stain (Sigma, Cat No. 18909) for 5-10 min followed by imaging at room temperature. 3D images consisting of 26 XY images with a Z-slice spacing of 0.204 µm (total Z-stack thickness 5.1 µm) were acquired at 100× magnification using a fluorescence microscope (Axioskop 2 MOT plus, Carl Zeiss, Inc.) equipped with a Plan Apochromat 100×/1.4 Oil DIC objective, an Axio Cam HRm camera and an HBO 100 Mercury lamp. Identical exposure times (50 ms) were used to acquire the green channel images whereas the exposure time for blue channel was adjusted for individual images based on the intensity of calcofluor staining.

For manual counting, images were deconvolved with theoretically generated PSFs using Axiovision software V4.9.1 SP2 followed by the generation of maximum intensity Z-projections (MIP) of both green and blue channels. All the deconvolved MIP images from WT and mutant strains were blinded and labelled as ‘1-44’, and their grey levels were set to ‘best fit’ in the Axiovision software prior to providing them to two individuals who manually counted peroxisomes in individual cells using the ‘measure events’ tool in Axiovision. Unlike manual counting, no post-processing was performed on the raw images used as the input for *perox-per-cell*.

##### *Strain construction*

All strains used in this study (Table S1) were constructed using lithium acetate-PEG-mediated transformations. For this, the parental wild type (WT) strain (*BY4742*) or those lacking *PEX30*, *VPS1*, *PEX3* and *INP1* ORFs [obtained from the *MATα* deletion library (Giaever *et al.*, 2002)] were transformed with a linear construct expressing GFP with Enhanced Peroxisome Targeting Sequence [ePTS1 (DeLoache *et al.*, 2016)] at its 3′ end from the *TDH3* promoter and followed by the *PGK1* terminator along with the *natMX6* selection cassette and targeted after the STOP codon of the *UBC9* ORF to fluorescently tag peroxisomes. *INP2* was deleted in the WT strain expressing *GFP-ePTS1* using the *hphMX6* cassette flanked by ~600bp sequences upstream and downstream of the *INP2* ORF.

**Table S1: Strain list**

| Name | Genotype | Alias |
|---|---|---|
| PPC1 | *MATα*, *ubc9::UBC9* (*natMX6*)-$P_{TDH3}$-*GFP-ePTS1*, *his3Δ1*, *leu2Δ0*, *lys2Δ0*, *ura3Δ0* | *BY4742*, GFPePTS1 |
| PPC2 | *MATα*, *ubc9::UBC9* (*natMX6*)-$P_{TDH3}$-*GFP-ePTS1*, *pex30Δ::KanMX4*, *his3Δ1*, *leu2Δ0*, *lys2Δ0*, *ura3Δ0* | *pex30Δ*, GFPePTS1 |
| PPC3 | *MATα*, *ubc9::UBC9* (*natMX6*)-$P_{TDH3}$-GFP-ePTS1, *vps1Δ::KanMX4*, *his3Δ1*, *leu2Δ0*, *lys2Δ0*, *ura3Δ0* | *vps1Δ*, GFPePTS1 |
| PPC4 | *MATα*, *ubc9::UBC9* (*natMX6*)-$P_{TDH3}$-GFP-ePTS1, *inp1Δ::KanMX4*, *his3Δ1*, *leu2Δ0*, *lys2Δ0*, *ura3Δ0* | *inp1Δ*, GFPePTS1 |
| PPC5 | *MATα*, *ubc9::UBC9* (*natMX6*)-$P_{TDH3}$-GFP-ePTS1, *pex3Δ::KanMX4*, *his3Δ1*, *leu2Δ0*, *lys2Δ0*, *ura3Δ0* | *pex3Δ*, GFPePTS1 |
| PPC6 | *MATα*, *ubc9::UBC9* (*natMX6*)-$P_{TDH3}$-GFP-ePTS1, *inp2Δ::hph*MX6, *his3Δ1*, *leu2Δ0*, *lys2Δ0*, *ura3Δ0* | *inp2Δ*, GFPePTS1 |

*Statistical methods*

To choose microscopy images that best represented the number of peroxisomes per cell or peroxisomal areas among a group of images, we first pooled all *perox-per-cell* results from those images, then performed Wilcoxon rank-sum tests to compare each individual image's results to the pool. The image with the least significant *P*-value was selected as the most representative.

FDR-adjusted *P*-values were computed using the Benjamini-Hochberg method (Benjamini and Hochberg, 1995).

*References*


Benjamini,Y. and Hochberg,Y. (1995) Controlling the false discovery rate: A practical and powerful approach to multiple testing. *J. R. Stat. Soc. Ser. B*, **57**, 289–300.

DeLoache,W.C. *et al.* (2016) Towards repurposing the yeast peroxisome for compartmentalizing heterologous metabolic pathways. *Nat. Commun.*, **7**, 11152.

Giaever,G. *et al.* (2002) Functional profiling of the *Saccharomyces cerevisiae* genome. *Nature*, **418**, 387–391.

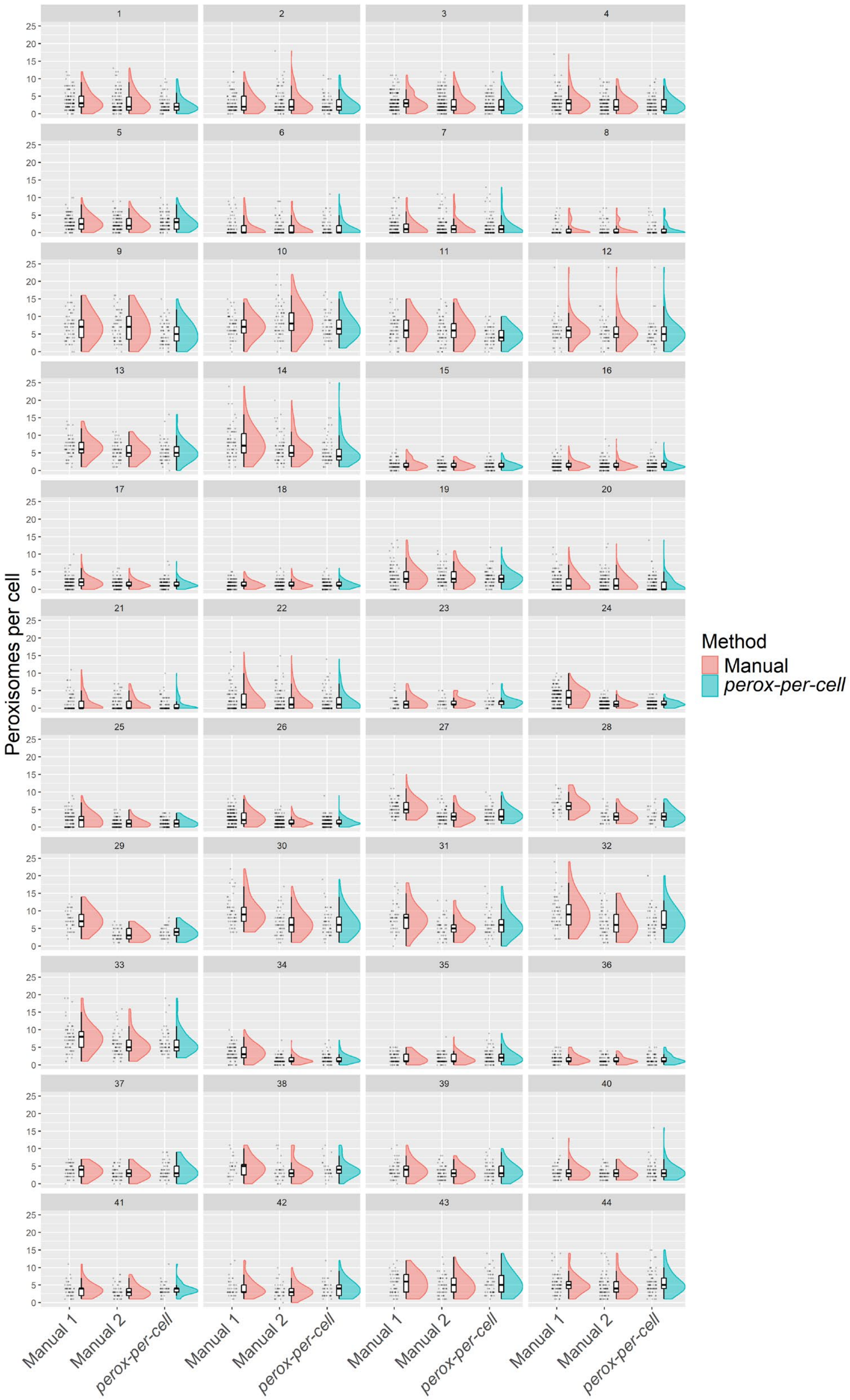


**Fig. S1. Distributions of per-cell peroxisome counts assessed manually or using *perox-per-cell* across 44 test images.** Adjacent jitter plots, box plots and half-violin plots are shown for each distribution. Images 6-8 are from the *inp2*Δ strain; 9-14, 30-33, and 43-44 are from *pex30*Δ; 15-18, 23-26, and 34-36 are from *vps1*Δ; 20-22 are from *inp1*Δ; the remaining images are from WT.

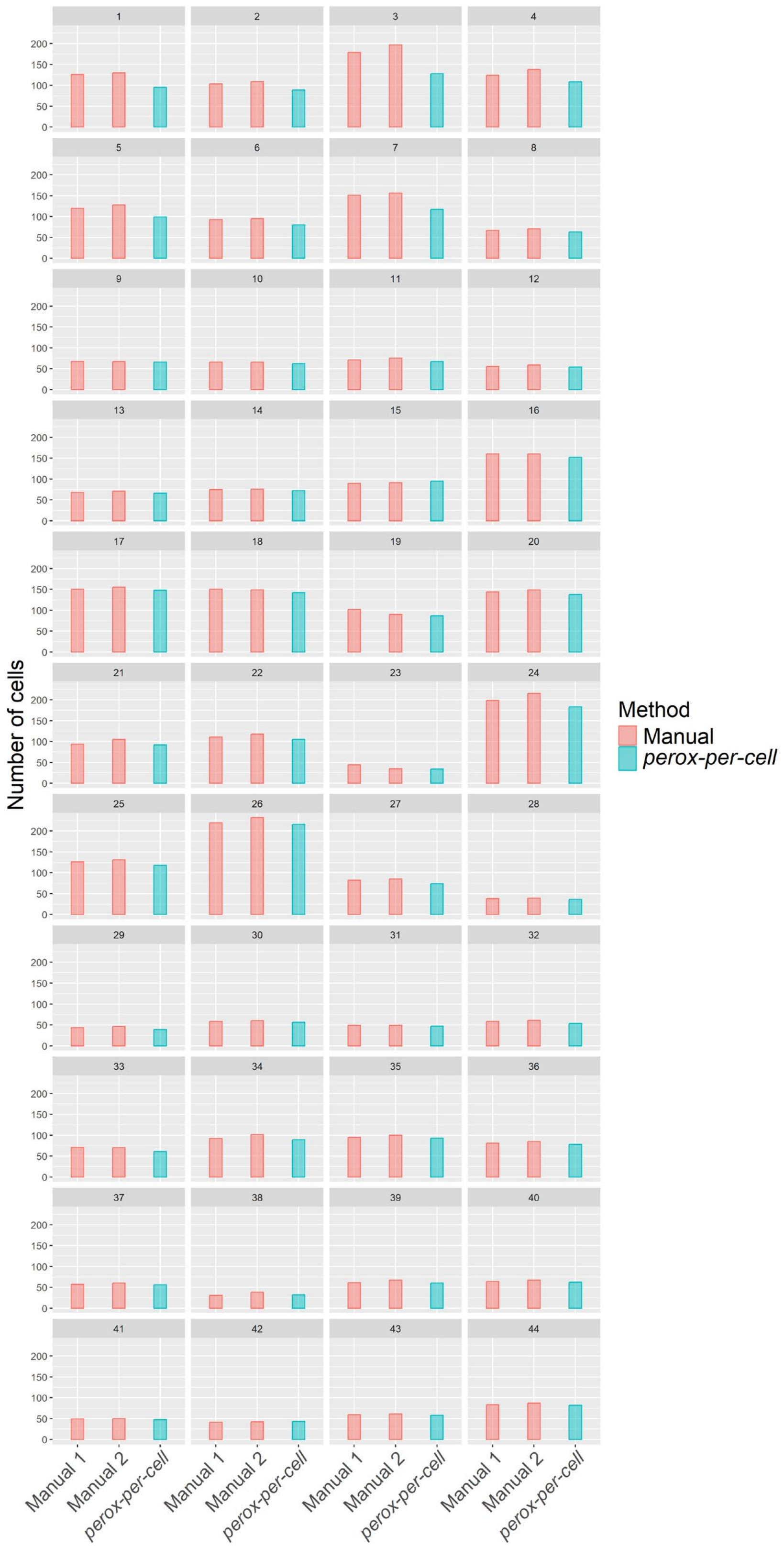


**Fig. S2. Number of cells counted either manually or using *perox-per-cell* across 44 test images.**
